# Behavioral variation in *Drosophila melanogaster*: no evidence for common alleles of large-effect at the *foraging gene* in a population from North Carolina, USA

**DOI:** 10.1101/004325

**Authors:** Thomas L. Turner, Christopher C. Giauque, Daniel R. Schrider, Andrew D. Kern

## Abstract

It has been postulated that natural populations of *Drosophila melanogaster* are comprised of two behavioral morphs termed "rover" and "sitter", and that this variation is caused mainly by large-effect alleles at a single locus. Contrary to common assertions, however, published support for the existence of common large effect alleles in nature is quite limited. To further investigate, we quantified the foraging behavior of 36 sequenced strains from a natural population, performed an association study, and described patterns of molecular evolution at the *foraging* locus. Though there was significant variation in foraging behavior among genotypes, this variation was continuously distributed and not significantly associated with genetic variation at the *foraging* gene. Patterns of molecular population genetic variation at this gene also provide no support for the hypothesis that *for* is a target of recent balancing selection. Though our data only apply to this specific population, we propose that additional data is required to support a hypothesis of common alleles of large effect on foraging behavior in nature.

## Introduction

IN 1980, Sokolowski [1] described a now classical difference in behavior between two mutant strains of *Drosophila melanogaster*. When placed in a food patch, larvae of one strain would stay there and eat (“sitters”) while the larvae of the other strain would crawl around while eating and visit other food patches (“rovers”). Further comparisons of a strain with the sitter trait and a strain with the rover trait provided evidence that differences between them could be explained mostly by variation at a single locus containing the *foraging* gene [2,3]. Though the original variation was described in strains that had been reared in the lab for decades, variation in foraging behavior was also found in natural populations, and a hypothesis was put forth that this behavioral variation is maintained by balancing selection in the wild [1,4,5]. In the decades since the original description of this variation, a considerable research effort has been focused on “reference” rover and sitter strains, thought to be representative of two natural morphs. These reference strains have been repeatedly compared and found to differ in many traits in addition to larval foraging, including adult movement patterns [6,7], energy storage (lipid vs. carbohydrates [8]), glucose homeostasis [9], thermotolerance [10], resistance to sleep deprivation [11], relative strength of short term vs. long term memory [12,13], use of "retroactive interference" [14], use of public vs. private information [15], and more. The picture painted is one in which *D. melanogaster* populations are composed of two coexisting behavioral “morphs”, differentiated in many phenotypic dimensions, that coexist through some type of balancing selection. The only comparable case we are aware of is variation at the *npr-1* gene in *Caenorhabditis elegans* [16,17] (though there may be non-recombining supergenes with diverse effects [18,19]). Similar to the *foraging* case in *D. melanogaster,* the *npr-1* allele from the N2 strain was found to cause solitary foraging, in contrast to the group foraging observed in other strains [20]. It has recently been postulated, however, that this large-effect allele arose as an adaptation to lab culture, and may not be maintained by selection in natural populations [21,22]. We feel that, based on published data, this same scenario cannot be ruled out in the case of variation at *foraging*. We first critique the existing evidence supporting the "common alleles of large effect" hypothesis, and then present additional data which fails to support this hypothesis. Inference from these new data are limited in scope, as we have only investigated a few dozen lines from a single population. Nonetheless, we feel it is important to publish these negative results so that a more complete picture of foraging variation may eventually be obtained.

### Previous Evidence for Bimodal Trait Distributions in Natural Populations

If the distribution of trait variation is bimodal, this supports a hypothesis that there is a factor of large effect involved (genetic or environmental). In the case of foraging behavior in *D. melanogaster,* several bimodal distributions have been reported. An impressively bimodal distribution is shown in a highly cited review that features the *foraging* story [23]. The source of these data is not reported in the review, but the distribution appears identical to figure 1 from de Belle, Hilliker and Sokolowski [24]. If this is indeed the source of these data, this bimodality does not address whether there is a common factor of large effect in natural populations. The bimodal distribution in de Belle, Hilliker and Sokolowski [24] is a comparison of only two "reference" strains: the EE (*ebony^11^*) strain (one mode) and the B15 strain (the other mode; see below for a further description of these genotypes). The variance around each mode is variation among replicate individuals of the same genotype.

**Figure 1.**
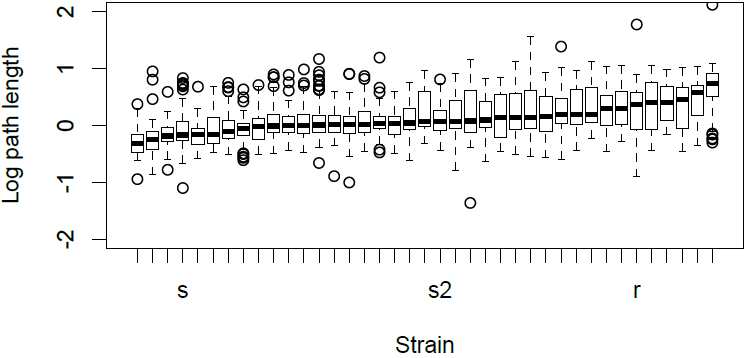
**Variation in larval foraging behavior**. Boxplot of foraging path lengths for 36 inbred RAL strains (not individually labeled, for clarity) and the reference rover (r), sitter (s) and sitter 2 (s2) genotypes. The median for each line is shown, surrounded by a box (1st to 3rd quartiles), whiskers (range excepting outliers), and circles (outliers calculated as more than 1.5 the interquartile range from the box); sample sizes per line vary from 28 to 81 (median n=54).

A bimodal distribution is also reported in a note published by the Drosophila Information Service [25]. This distribution shows variation in foraging behavior in larvae collected from fallen pears near Toronto. The distribution of foraging path lengths for these larvae is bimodal, with one mode between 60–80 mm (named "rovers") and the other mode at 0-20 mm (named "sitters"). The bimodality of this distribution supports a hypothesis that there is a factor of large effect in the data, but this factor is not necessarily genetic. These larvae were sampled from nature and assessed directly, so variation among them could be due to genes, the environment, and/or gene-by-environment interactions. Indeed, major effects of environmental factors were later quantified in laboratory assays [26], and it was later noted that "a carefully controlled environment is required to minimize the phenotypic overlap between distinct genotypes" [24]. It therefore seems plausible that the alternative possibility—individuals of the same genotype behaving differently due to environmental influence—cannot be excluded. Indeed, it was later stated that "flies with rover alleles can be made to behave as sitters after a short period of food deprivation" [23].

Evidence for a genetic effect on foraging distance was reported in the original foraging paper, but not for genotypes recently sampled from nature [1]. Larvae of one strain, carrying a *white^blood^* mutation affecting eye color, traversed a much larger area when feeding ("foraging") than the other strain, which carried a mutant *ebony^11^* allele affecting body color. This behavioral difference did not seem to be caused by either of the pigmentation mutations, as *ebony* is on the third chromosome, *white* is on the X, and the behavioral difference mapped almost entirely to the second chromosome. Individuals with the second chromosome from the *ebony^11^* strain were termed "sitters", and those with the *white^blood^* second chromosome were termed "rovers". Though given the same names as the larvae sampled from nature, no evidence is reported that supports this inference. Differences between these lab stocks could have arisen during lab culture. Flybase.org lists the first reference to the *ebony^11^* allele as a paper from 1926 [27]: 54 years of lab culture is likely equivalent to over 1000 generations since this strain was sampled from nature. The earliest reference we could find to the *white^blood^* strain was published in 1944 [28], which is also a considerable interval.

The strongest evidence for the existence of a common allele of major effect in nature, to the best of our knowledge, was presented by Sokolowski and colleagues in 1997 [5]. An outbred laboratory population was founded from 500 flies, again collected from a Toronto orchard, and 500 individual larvae were assayed from this population after a year of lab culture. The trait distribution of these individuals was once again bimodal [5], and appears very similar to the distribution of larvae measured directly from nature [1,25]. Moreover, the differences between "rover” and "sitter" larvae from this population failed to complement the sitter mutation from the *ebony^11^* lab strain, suggesting that there was variation at the same locus. Together, these data do support a hypothesis of common, large-effect alleles in nature. However, these data on their own still allow room for doubt, especially in light of the unprecedented nature of the foraging story. Possible critiques of these data include the complications of epistasis when performing complementation tests in uncontrolled genetic backgrounds [29]. In addition, the genotypes found to be bimodal were not isolated independently from nature. Instead, a large outbred population was adapted to the lab for a year, then individuals from this population were assayed. This means that allele frequencies could evolve between the sampling and measurement, so that a rare sitter allele (for example) could become more common. Indeed, the same paper proposes that these alleles can rapidly change in frequency in lab culture due to density-dependent selection. A more accurate procedure for investigating the frequency of natural genotypes in *Drosophila* is to sample individual females and use them to found independent isofemale or inbred lines. This method was used by Baur and Sokolowski [4], who show little support for the common alleles of large effect hypothesis. Fifteen fertilized females were collected from nature (source population not stated), and an isofemale line was started from each female’s descendants. Path lengths were found to vary significantly among these 15 lines, implicating heritable (likely genetic) variation. However, the trait values of all lines shown would lead to their classification as rovers based on the previous definition. The data presented appear to vary continuously from ~50-85 mm, without two modes [4]. Thirteen of these lines were not classified as either rover or sitter, but the line (B1) with the shortest path length was classified as a sitter. This appears arbitrary, as the behavior of this line appears more similar to the "roving" *white^blood^* stock than to the "sitting" *ebony^11^* stock [1]. The strain with the longest path length (B15) was designated a "rover". The B1 and B15 lines were referred to as "extreme", but they do not appear to be statistical outliers compared to the other 13 lines [4]. The primary support for a hypothesis that the genetic difference between B1 and B15 was related to the difference between *white^blood^* and *ebony^11^* was that the mutation(s) causing the difference between the B1 and B15 strains were later mapped mainly to the second chromosome [4]. This is consistent with a large-effect allele, but it should be noted that the second chromosome comprises ~2/5 of the euchromatic *D. melanogaster* genome, and could therefore harbor causal variants in many genes [30]. The strain with the longest path (B15) appears to be the strain that was later used as the "reference" rover strain, renamed the *for^R^* strain (sometimes also referred to as R, B15B15 or BB). The strain with the shortest path, however, does not appear to be the reference sitter strain (EE, S, or *for^s^)* that was used in subsequent mapping experiments. Instead, strains with the second chromosome from B15 were compared to strains with the second chromosome from the *ebony^11^* lab stock in later work (see below).

Foraging behavior has more recently been reported for a second set of stable genotypes, 22 inbred lines, and these lines again varied continuously with a single mode [31]. These lines were all referred to as "sitter-like", though we would suggest that they do not support a binary rover vs. sitter classification scheme. Other genotypes have also been compared, but these cases do not support the common allele of large effect hypothesis either. For example, artificial selection was performed on a *sepia* mutant stock collected by Timothy Prout near the University of California, Riverside, and kept in the lab for 15 years before the start of the experiment [32]. The trait values of this stock before selection would classify them as "rovers" based on the previous definition [1,25], as the average path lengths were between 60 and 110 mm [32,33]. Selection was successful in both directions, indicating that there was heritable variation in path length in this stock [33]. However, this is not evidence that the stock was segregating a "sitter" allele of large effect. After selection, in fact, the population selected to be sitter-like would still have qualified as "rovers" based on the definitions provided [33]. These data are therefore consistent with multiple alleles of small or moderate effect.

Additional collections were made from the Toronto pear orchard in 1986, and laboratory stocks collected from different regions of a single pear were found to differ in average path length [34]. Only the mean values are shown, however, and no evidence of a bimodal distribution is presented. Stocks with the second and third chromosomes from the *ebony^11^* strain (now called EE) and *white^blood^* (now WW) stocks (and stocks derived from them) were reassayed for comparison to this fresh collection, and the newly sampled short-path-length strains had intermediate trait values compared to the EE and WW lab strains [34]. The shorter path strains were called sitters, and the longer path strains were called rovers, but this appears arbitrary compared to previous definitions. The only additional report of a bimodal distribution we are aware of is for a trait correlated with larval foraging distance, but this appears to be a truncation artifact, as flies given a larger area displayed continuous variation with a single mode [35]. Nearly all other work on rover and sitter that we are aware of focused only on the "reference" strains, discussed further below, and therefore cannot speak to the frequency of *foraging* alleles in nature. We therefore feel that these data, though they raise the interesting possibility of common behavioral morphs in nature, are insufficient to strongly support such a hypothesis.

### Published Data Connecting Genotype and Phenotype

Efforts to connect genotype and phenotype have focused nearly exclusively on three reference strains: *for^r^, for^s^*, and *for^s2^*. This is understandable, due to the labor-intensive nature of behavioral characterization, but limits inference regarding natural populations. First, it is not clear that the *for^s^* strain used has an allele recently isolated from nature, as is sometimes asserted [7,8]. Efforts to map the genetic basis of path length to a sub-chromosomal level were first published in de Belle, Hilliker, and Sokolowski [24]: the sitter allele used in these efforts is the one isolated from the *ebony^11^* lab stock. The sitter strain, before being renamed*for^s^,* is described as "EE". Based on [1] and [34], this strain seems to have the second and third chromosomes, and therefore the *foraging* allele, from the *ebony^11^* stock. The rover strain, renamed *for^R^*, is also referred to as B15B15, and therefore appears to have the second and third chromosomes from the B15 strain [4]. This B15 or R stock is referred to as *for^R^* by de Belle, Sokolowski, and Hilliker [36], and appears to have become the reference rover strain. Supporting this inference is the common reference to the *for^R^* strain as the parent of the irradiated strain *for^s2^,* produced from B15B15 by [24]. However, de Belle, Sokolowski, and Hilliker [36] also clearly indicate that the strain renamed *for^s^* is the EE strain, which has the *foraging* allele from the *ebony^11^* stock. The final identification of the gene *dg2* (a cGMP-dependent protein kinase, subsequently renamed *foraging)* as the gene harboring the *for^R^* and *for^s^* alleles does not state the source of the *for^s^* allele, but the *ebony^11^* stock seems likely as it was a continuation of the earlier mapping work, and uses the same stock name [2]. Other sitter-like chromosomes that failed to compliment the *ebony^11^* allele have also been referred to as *for^s^* in the literature [5], so it is possible that other alleles were used.

Regardless of the source of the *for^s^* allele, the *for^R^* and *for^s^* strains were created by chromosome extraction [1,34,36], and therefore differ at a very large number of loci. As a result, it is not possible to associate traits with *foraging* alleles simply by measuring the phenotype of these strains (e.g. [15,37]). Thankfully, most studies also use the *for^s2^* strain [8–10,12,38]. This sitter-like strain was created from the reference rover strain using high doses of gamma radiation (5000 rad; [24]). This strain therefore shares a common background with *for^R^*, though it seems likely to have secondary mutations, despite assertions that it differs from *for^R^* only at *foraging* [7,14,38]. Some trait associations have additional support from transgenic manipulation [7,11,12,38], including the original association between *foraging* and larval path length [2]. To our knowledge, however, the genome sequences of the reference strains have not been reported. It has also come to our attention that one of the four lines of evidence used to link the *foraging* gene to larval path length was later retracted in a personal communication to Flybase.org [3]. A strain thought to have a P-element insertion in *foraging* was determined to be a sitter strain, and excision of the element caused a reversion to rover-like behavior [2]. Upon reanalysis it has apparently been determined that this insertion is in the gene *lilliputain* rather than *foraging* [3].

It therefore seems that, although variation at the gene *foraging* can likely affect interesting behaviors, additional evidence is required to support a hypothesis of large-effect alleles at high frequency in nature, which affect many aspects of physiology and behavior, and thereby create two behavioral "morphs" maintained by balancing selection. To collect additional data, we have quantified larval foraging behavior on a set of inbred lines isolated from a wild population and attempted to associate trait variation with genotype. Our analysis fails to support the common-alleles of large effect hypothesis in that we were unable to associate variation at the *for* locus with foraging behavior. Population genetic analyses of variation at *for*, relative to the rest of the genome, also do not support a hypothesis that *foraging* is a target of strong, recent balancing selection in the *Drosophila* genome. As we have only investigated a few dozen lines from a single population, these negative results should only be taken as one small piece of a larger, and still unresolved, puzzle.

## Results

Foraging path lengths were measured in the reference rover and sitter strains as well as 36 natural isolates of *D. melanogaster* from the RAL collection (figure 1). These strains were started from females captured in Raleigh, North Carolina, and underwent full-sib mating for 20 generations [39]. This procedure produces an inbred genotype while allowing very little opportunity for selection (though some alleles, like recessive lethals, are purged).

When measured by us, the *for^s^* and *for^s2^* strains were found to have shorter average path lengths (1.30 and 1.71 cm) than the *for^R^* strain (2.66 cm), as expected. The foraging paths of the *for^s^* strain were significantly shorter than the *for^s2^* strain (t-test p=0.002), and both are significantly shorter than the *for^R^* strain (t-test p=1.8E-6 and p=0.016, respectively). All reference strains traveled shorter distances in our assay compared to published data. One recent publication reported *for^s2^* larvae travel approximately 3.5 cm and *for^R^* travel approximately 7 cm [31]; other publications report even larger values, such as 10.89 cm and 17.83 cm for *for^s^* and *for^R^*, respectively [6]. Note that the *for^R^* strain is reported to travel approximately twice as far as *for^s^* in both cases, which agrees with our data.

Data from the 36 inbred RAL strains are distributed with only one mode (figure S1). The mean path length among lines is significantly non-normal (Shapiro-Wilk’s Test p<0.001), but not after log_10_ transformation (Shapiro-Wilk’s Test p=0.075). Only three strains had shorter paths than the *for^s^* strain, while only five strains traveled farther than the *for^r^* strain: the majority of strains fell between these two reference points (figure 1). We estimate the proportion of phenotypic variation resulting from genotypic variation, or the broad-sense heritability, as 20% (estimated as the variance explained by line).

To investigate whether any of the phenotypic variation among RAL strains may be due to alleles at the *foraging (for)* gene, we performed an association study. Though genome-wide variation data is available for these lines, we limited our analysis to the *for* locus because the *a priori* expectation of an association at this locus reduces the loss of power from multiple testing. As annotated at flybase.org (genome assembly version 5.54) the *for* gene is found on the left arm of chromosome 2 from base pairs 3,622,074-3,656,953; 5,000 bp to either side were also included as potential regulatory regions. Association tests were performed for each of 470 variants (single nucleotide polymorphisms (SNPs) and small insertion-deletion (indel) mutations) that are 1) mapped to this interval, 2) biallelic, and 3) present in at least 5 lines included in our analysis (~15% allele frequency or greater). The most significant association in this interval had a p-value of only 0.022, uncorrected for multiple tests (see supplement for full results). With 470 parallel tests, this variant is not close to the Bonferroni-corrected significance value of 8.24E-5.

We used a power simulation to explore the possibility that one of the variants we tested has a large effect on our phenotype. The hypothesis we are trying to reject is that there is a variant of large effect among the 470 variants tested. Large effect, in this context, is relative to the reference *for^r^* and *for^s^* strains. If the phenotypic difference between reference strains is due mainly to one variant, simulations indicate that this variant was not among those tested (figure 2). If the *for^r^* allele was at 50% frequency in our sample (consistent with the postulated 70/30 split between dominant rover and recessive sitter phenotypes), and explained at least 60% of the phenotypic difference between *for^r^* and *for^s^*, there is a 98% probability that it would have p<0.022 (the lowest p-value observed). This probability drops to 95% if the *for^r^* allele is at 30% frequency in our sample, and 72% if it is at only 20% frequency. In contrast, an allele explaining only 20% of the phenotypic difference between reference strains would likely remain undetected in a sample as small as ours (figure 2).

**Figure 2.**
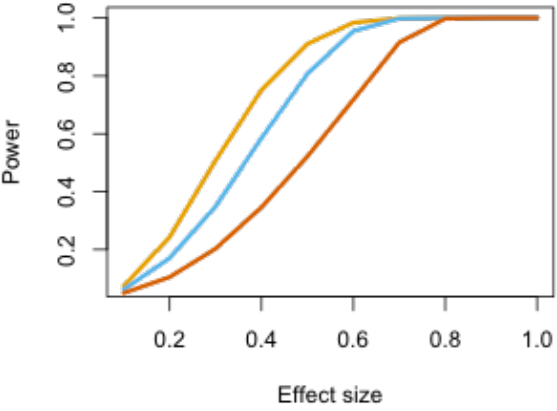
**Power analysis**. Simulated data were used to asses the power of our analysis to reject a hypothesis of common, large-effect alleles. Large-effect in this case is relative to the difference between reference rover and sitter strains. An x-axis value of 1.0 would be an effect equal to the difference between strains, while an effect size of 0.2 is 20% of the difference between strains. Power indicates the proportion of simulations with p-values lower than any observed in our data. Three allele frequencies were simulated: 50% (yellow), 30% (blue), and 20% (red).

### Patterns of Polymorphism and Divergence

To investigate whether there is any signature of balancing or frequency dependent selection acting at the *for* locus [40], we examined patterns of polymorphism and divergence in the *Drosophila* genome. This analysis utilized the full 154 strains in the Drosophila Genetic Reference Panel (DGRP [41]) that were sequenced with Illumina technology (Methods)—this set includes the majority of the strains used in our association study. If some flavor of balancing selection was acting at this locus we would expect to see elevations in polymorphism relative to divergence either throughout the locus or in a particular region of the locus that is the target of such selection. The *for* locus, which encodes a rather long protein, has 18 nonsynonymous polymorphisms and 76 synonymous polymorphisms in the DGRP sample. Accounting for the total length of the longest transcript at *for* this amounts to 0.0063 nonsynonymous sites per base pair (55^th^ percentile among genes the recombining region of chromosome arm 2L whose protein lengths are ≥ that of *for*) and 0.028 synonymous sites per base pair (89^th^ percentile). Nucleotide diversity paints a rather similar picture. Locus-wide π=0.0072, which is greater than median diversity but only at the 82^nd^ percentile among genes. Nucleotide diversity at synonymous and nonsynonymous sites appears no different; π_N_=0.00061 (61^st^ percentile on 2L) and π_S_=0.017 (89^th^ percentile). Thus while *for* shows above average diversity, it does not seem to be a strong outlier as might be expected from a balanced polymorphism.

To formally test the neutral model at the *for* locus we looked at the site frequency spectrum of polymorphism and comparisons of polymorphism and divergence. Under a model of balancing selection one expects to see a skew in the site frequency spectrum towards intermediate frequency polymorphisms. Using Tajima’s *D* statistic [42], this would translate into a strongly positive value of *D*. At *for* we find that *D* = 0.6074, which when tested against coalescent simulations under the standard neutral model yields a p-value of *p* = 0.204 in a one-side test. Further, compared to the empirical distribution of *D* statistics for protein coding genes on 2L, *for* is at the 75^th^ percentile. This observation suggests that the site frequency spectrum at *for* is not particularly unusual or strongly skewed towards intermediate frequencies. Sliding windows of Tajima’s *D* (Figure 3) do show at least one interesting window of positive *D* (i.e. intermediate frequency polymorphisms) near the 3’ end of the coding region, with a value of *D* = 1.25. From the empirical distribution of *D* throughout chromosome 2L this window appears to be an outlier at the 0.05 level.

**Figure 3.**
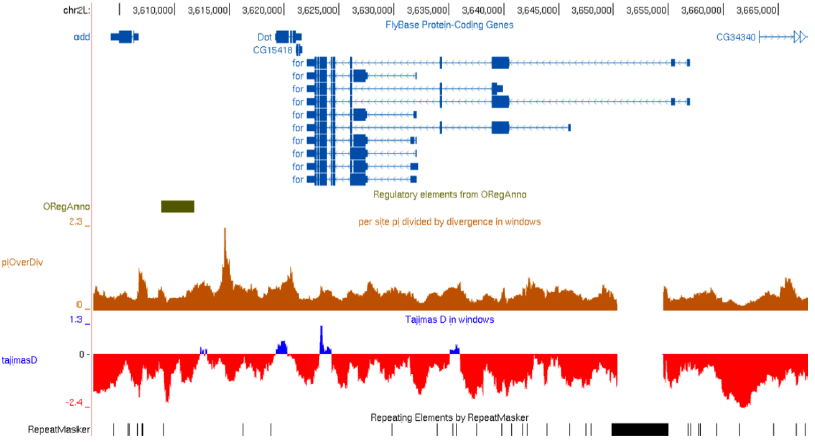
**Polymorphism and divergence at the *for* locus**. An image from the UCSC Genome Browser (Kent et al. 2002) showing the various isoforms of for and its flanking regions. The ratio of nucleotide diversity to polarized divergence on the *D. melanogaster* lineage and values of Tajima’s Dare shown in 1 kb windows sliding every 100 bp, labeled as piOverDiv and tajimasD, respectively. Regulatory elements from ORegAnno (Montgomery et al. 2006; Griffith et al. 2008) are also shown, as are repetitive elements from RepeatMasker (http://www.repeatmasker.org). The large masked element near the 5’ end of the gene is a Copia insertion that varies in copy number within the DGRP population sample.

Another strong prediction under a model of balancing selection is that the ratio of polymorphism and divergence should be skewed in favor of too much polymorphism relative to divergence under neutrality. We tested this prediction in three different ways. First we used an unpolarized McDonald-Kreitman (MK) test to examine patterns of polymorphism and divergence at synonymous and nonsynonymous sites at *for* using *D. simulans* as an outgroup. The unpolarized MK test (Table 1) is non-significant (*p*=0.30; Fisher’s exact test) and has a Neutrality Index (N.I.) of 0.64 [43]. Comparing the N. I. of*for* to the empirical distribution from protein-coding genes on 2L, we find that *for* is near the middle of the distribution (28^th^ percentile). The second test of polymorphism and divergence we performed was a polarized MK test where we examined only fixations along the *D. melanogaster* lineage. The results are qualitatively similar in this case (*p*=0.26; N.I.=0.53; 8^th^ percentile). Thus neither polarized nor unpolarized MK tests can reject the neutral model. We obtain similar results when using variant calls from the complete set of sequenced lines (including those sequenced by Roche 454 as well as Illumina) from freeze 2 of the DGRP data set (ftp://ftp.hgsc.bcm.edu/DGRP/freeze2Feb2013/), as well as a sample of African genomes.

**Table I.**
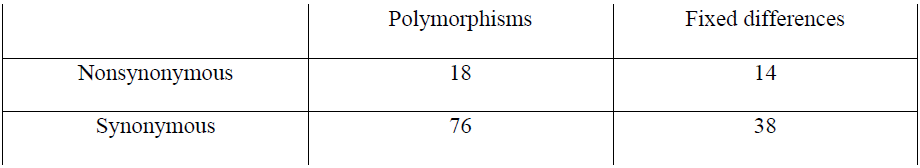
**Unpolarized McDonald-Kreitman test of *for*.**

Finally we were interested in examining the ratio of polymorphism and divergence directly from the entire locus and from individual windows, including upstream and downstream regions that may contain important regulatory elements, in a Hudson-Kreitman-Aguadé like test [44]. The ratio of nucleotide diversity to divergence (again using only fixations along the *D. melanogaster* lineage) for the whole *for* locus is equal to π/div=0.419. Comparing this ratio to that of genes on 2L encoding a protein at least as large as *for’s, for* is on the higher-end of the empirical distribution but not a strong outlier (89^th^ percentile). We also examined sliding windows of the ratio of polymorphism to divergence (π/div) and the results from this analysis are shown in figure 3. This analysis revealed perhaps the only remarkable feature of genetic variation in the *for* region. While the ratio of polymorphism to divergence throughout the locus is below one and rather average for the genome, there is a strong peak in polymorphism relative to divergence approximately 8kb 3’ of the *for* locus. Unfortunately this peak does not overlap any known regulatory sequences of *for* and moreover there are two proteins coding genes between this peak and the end of the *for* coding sequence. Moreover, the interesting window identified by Tajima’s *D* above does not stand out as an outlier with respect to patterns of polymorphism and divergence. Thus while this may be an intriguing finding we have no ability to say whether this is associated with *for* or not. After observing that this region was unusual, we performed an association to see if any variants in this interval were associated with foraging behavior. Thirty-eight SNPs in this interval (3,613,450 - 3,613,450) were included in the GWAS. The most significant association of these is a G/T polymorphism at 6,614,038 (p=0.0047; p = 0.16 after correcting for 38 parallel tests).

It is also possible that structural polymorphisms at the *for* locus contribute to variation in foraging behavior. We therefore asked whether any deletions or tandem duplications detected by Zichner et al. [45] in a sample of 39 of the DGRP genomes were found overlapping or within 10 kb of *for*. Two small deletions (70 and 124 bp), both found in 4 of these 39 genomes, are located within an intron of and upstream of *for*, respectively. The Copia LTR insertion shown in figure 3 also varies in copy number, being present in only 3 genomes. We cannot rule out the possibility of these variants affecting foraging behavior—if they are not in linkage disequilibrium with any of the SNPs or indels included in our association study then we are unable to detect any effect they may have. No tandem duplications detected by Zichner et al. appear within 10 kb of *for*.

## Discussion

There is little doubt that decades of work on variation at the gene *foraging* has proven interesting. Considerable mapping efforts culminated in the connection of variation in larval behavior to the gene *foraging* [2], and this catalyzed additional research into this interesting gene in other systems [46], The prevalence of large-effect alleles in nature is not well established, however, despite published claims that *D. melanogaster* populations are comprised of 70% rovers and 30% sitters, and that these behaviors are caused mainly by variation at *foraging* [2,23]. Published data are subject to alternative interpretation due to uncontrolled environmental variation [1,25] or mass culture [4]. The only published data using independently sampled genotypes reared in a common environment, that we are aware of, does not appear to support a gene of large effect [4,31]. To gather additional data about variation in larval foraging behavior, we have assayed a set of 36 inbred lines derived from flies collected in the Raleigh, North Carolina farmer’s market. The distribution of larval foraging behavior among these lines is not bimodal, nor is trait variation associated with genetic variation at or near the *foraging* locus.

Using standard analyses from molecular population genetics we sought to detect any signature of balancing selection that may maintain two (or more) alleles at *for* in nature as has been suggested in the literature [40]. None of our analyses reveal evidence for balancing selection at *for*, nor do they suggest that patterns of variation at *for* are somehow unusual within the *D. melanogaster* genome. Given the very large sample of genomes in this analysis, we have no reason to believe that our statistical tests should be in any way underpowered to detect recent strong balancing selection. Thus, from our failure to reject the null model of neutrality for both patterns of polymorphism and divergence as well as the site frequency spectrum, we conclude that strong balancing selection at *for* beginning recently (i.e. following migration out of Africa) is unlikely. However, more ancient balancing selection, allowing more time for recombination to whittle down the association between flanking markers and the site of selection, would be more challenging to detect. That said we find no significant elevation of the ratio of polymorphism to divergence even on a local scale with the protein coding locus itself.

The negative results of our association study come with major caveats. One possibility is that the trait we measured is not a high-fidelity replication of the "foraging behavior" measured in previous publications. The *for^R^* strain had a path length about twice as long as *for^s^* in our data, consistent with published reports [31]. The lengths of all paths in our data are shorter than reported for these strains previously, however, despite careful replication of the phenotyping protocols published earlier. If subtle differences in larval condition or assay environment are crucial and different, the effect of *for* might fail to be expressed. Furthermore, we only assessed larval foraging distance in the presence of food. This is consistent with and directly comparable to most published data, including both published bimodal distributions. It seems possible that *foraging* variation would be more apparent if the difference in path length with and without food was assayed—especially if there were other genetic variation affecting overall motility. In contrast to the isofemale lines and outbred populations described previously, the RAL lines used here are inbred, and any recessive variation in overall performance could be a more substantial confounder. We also note that there could be genetic variants at the *for* locus that are not annotated in the RAL genome alignments or have not been uncovered by studies of structural variation.

Large-effect mutations have been found in nature for traits like pigmentation and insecticide resistance, but it is unclear how often such large effects should be expected [47,48], especially given that in theory such variation in fitness related traits should be rare. Few natural alleles of large effect have been found for behaviors, and the only other *Drosophila* case, the effect of the period gene on courtship song evolution [49], has now been shown to be an artifact [50]. Common alleles of large-effect at *foraging* would therefore be an excellent opportunity to investigate the molecular architecture of natural variation in behavior, and we hope that our negative result will motivate further work addressing this hypothesis.

## Materials and Methods

### Fly strains

The reference *for^r^, for^s^* and *for^s2^* strains were provided by Marla Sokolowski. We also used inbred lines from the Raleigh (RAL) collection, provided by the Bloomington Drosophila Stock Center (these lines are part of the Drosophila Genomic Reference Panel, or DGRP). These lines were collected by the Mackay lab in Raleigh, North Carolina, and each line underwent full-sibling inbreeding for 20 generations to eliminate most genetic variation [39,41]. All lines were maintained in 25×95 mm vials on molasses medium in standard *Drosophila* incubators, at 25°C under a 12-hr light/dark cycle.

### Behavioral measurement

The rover/sitter phenotype was measured for each line similarly to previously published methods [51]. Oviposition bottles were prepared with grape juice agar plates with yeast paste added. Parents were allowed to lay eggs freely for 1 hour at 25°C, after which the plates were incubated overnight at 25°C. Twenty-four hours later, individual newly hatched larvae were carefully removed from the oviposition plates using forceps and transferred to Petri plates containing 35 mL standard *Drosophila* food. No more than 50 larvae were grown in each plate to avoid overcrowding. They were allowed to grow in these plates until 96 hours post-hatching. At this time, each third instar larva was removed from the food and individually tested for foraging behavior. Only larvae found within the food were tested. Any larvae on the food surface or on the surface of the Petri plate were not used. For the behavioral assay, a thin layer of yeast paste was applied to a custom-built plastic plate using a rubber squeegee. A larva was placed into the middle of the yeast paste, the yeast paste was covered with half a Petri plate, and the larva moved freely for five minutes. After this time, the larval trail, visible in the yeast paste, was traced by pen onto the covering Petri plate. The marked plate was photographed, and the length of the larval path was measured using the ImageJ software package. Sample sizes per line varied from 28-81 (median n=54); raw data are available as supplementary data. The R software package was used to further analyze the results.

### Association study

Associations were conducted using the log_10_ transformed median path length for each line. A catalog of genetic variation was obtained on 10/13/2014 from "freeze 2" data available at https://www.hgsc.bcm.edu/arthropods/drosophila-genetic-reference-panel as a "binned" VCF file. Data from two of the 36 lines phenotyped, RAL-765 and RAL-514, were not omitted from the available data and therefore not included in our association study. For the remaining 34 lines, we tested all variants that 1) were between base pairs 3,622,074-3,656,953 on the left arm of chromosome 2, which includes the *for* gene and 5,000 base pairs to either side, 2) were biallelic, and 3) were present in at least 5 of our 34 lines. A t-test was used to test for association at each variant, excluding any strain annotated as a heterozygote. Full results are available as supplementary data.

We explored statistical power at a variety of allele frequencies and effect sizes using a simulation. For a given allele frequency and effect, we sampled without replacement from the distribution of line phenotypes. We then added an effect to these sampled averages and tested them vs. the other line averages. For example: we start with a vector of trait values for the 34 lines with genotypes and phenotypes. For a simulated allele at 20% frequency, we randomly sample 20% of these trait values and each is increased by the effect size in question. We then log transform these values and use a t-test is used to compare the 20% modified data to the 80% unmodified data. We did this 10,000 times for each combination of allele frequency and effect size, and determined what proportion of these tests had lower p-values than any of the 470 observed p-values in our association study. If, for example, 80% of the simulations for a given combination of allele frequency and effect size had lower p-values than any observed in our real data, this indicates that there is a 20% chance that one of our tested alleles at that frequency has such an effect.

### Molecular Population Genetics

All population genetic analysis was performed using the set of Illumina-sequenced lines from Mackay et al. [41]. To test for selection acting at the *for* locus we performed a lineage-specific McDonald-Kreitman test [52] and compared results from that test to lineage-specific McDonald-Kreitman tests from every other protein coding locus throughout the genome using *D. simulans w501* and *D. yakuba* genomes [53] as outgroups for polarization of fixed differences. A similar test was also performed using unpolarized comparisons. For summary statistics of patterns of polymorphism and divergence we calculated nucleotide diversity [54], Tajima’s *D* [42], and the ratio of nucleotide diversity to divergence on the *D. melanogaster* lineage ignoring repetitive regions masked by RepeatMasker (repeatmasker.org). These statistics were also calculated on synonymous and nonsynonymous sites separately. For nearly every summary statistic, we calculated the same statistic for the longest transcript (examining coding sequence only) from each protein coding locus throughout chromosome arm 2L (on which for resides), to establish the genome-wide empirical distribution from which to compare population genetic summaries at *for*. We limited this analysis to genes whose longest transcript was at least as long as that of *for*, to prevent greater variance at smaller genes from pushing *for* toward the center of the distribution. This analysis only included genes on the recombining portion of 2L (which we defined as positions 1,000,000-17,000,000 after examining data from Comeron et al.). We also compared *for* to all genes genome-wide and obtained highly concordant results (data not shown). All population genetic analysis was performed using programs written in Ruby, Python, or C, and are available on GitHub (https://github.com/kern-lab/Turner_et_al_for), as are the steps we used to perform alignments and format polymorphism data for use by these scripts..

## Acknowledgements

We are grateful to the Sokoloswki lab for providing stocks and to the Mackay lab for constructing the RAL collection. We thank A. Shanku for preparing the alignments and variant calls used for the majority of population genetic analyses. Funding was provided by the National Institutes of Health (R01 GM098614 to TLT) and by support for to the Center for Scientific Computing at UCSB from the US National Science Foundation (NSF MRSEC DMR-1121053 and NSF CNS-0960316). ADK was funded in part by NSF MCB-1052148 and by DOE/USDA 124336. DRS was supported by the National Institutes of Health under the Ruth L. Kirschstein National Research Service Award F32 GM105231. Stocks obtained from the Bloomington Drosophila Stock Center (NIH P40OD018537) were used in this study.

---

**Figure S1.**
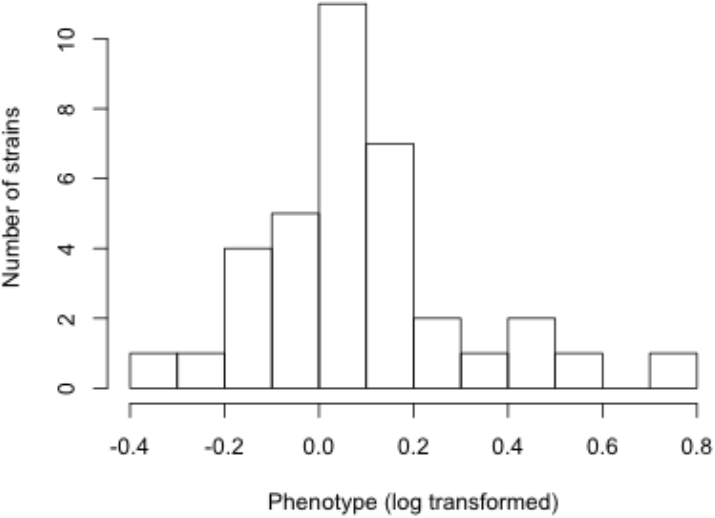
Histogram of variation in larval foraging behavior. Histogram of log transformed median foraging path lengths for 36 inbred RAL strains.

